# Estradiol promotes and progesterone reduces anxiety-like behavior produced by nicotine withdrawal in rats

**DOI:** 10.1101/842252

**Authors:** Rodolfo J. Flores, Bryan Cruz, Kevin P. Uribe, Victor L. Correa, Montserrat C. Arreguin, Luis M. Carcoba, Ian A. Mendez, Laura E. O’Dell

**Author notes:** Corresponding author: Laura E. O’Dell, Ph.D. Department of Psychology, The University of Texas at El Paso, 500 West University Avenue, El Paso, TX 79968.

## Abstract

The present study assessed sex differences and the role of ovarian hormones in the behavioral effects of nicotine withdrawal. *Study 1* compared physical signs, anxiety-like behavior, and corticosterone levels in male, intact female, and ovariectomized (OVX) female rats during nicotine withdrawal. Estradiol (E2) and progesterone levels were also assessed in intact females that were tested during different phases of the 4-day estrous cycle. *Study 2* assessed the role of ovarian hormones in withdrawal by comparing the same measures in OVX rats that received vehicle, E2, or E2+progesterone prior to testing. Briefly, rats received a sham surgery or an ovariectomy procedure. Fifteen days later, rats were prepared with a pump that delivered nicotine for 14 days. On the test day, rats received saline or the nicotinic receptor antagonist, mecamylamine to precipitate withdrawal. Physical signs and anxiety-like behavior were assessed on the elevated plus maze (EPM) and light-dark transfer (LDT) tests. During withdrawal, intact females displayed greater anxiety-like behavior and corticosterone levels as compared to male and OVX rats. Females tested in estrus (when E2 is relatively low) displayed less anxiety-like behavior and corticosterone versus all other phases. Anxiety-like behavior and corticosterone were positively correlated with E2 and negatively correlated with progesterone. Intact females displaying high E2/low progesterone displayed greater anxiety-like behavior and corticosterone as compared to females displaying low E2/high progesterone. Lastly, OVX-E2 rats displayed greater anxiety-like behavior than OVX-E2+progesterone rat. These data suggest that E2 promotes and progesterone reduces anxiety-like behavior produced by withdrawal.

## 1. Introduction

Clinical reports suggest that stress is a major factor that promotes tobacco use in women. For example, women report more often than men that they smoke to avoid negative affective states during abstinence (Stanton, 1995). Women also display stronger negative affective states, such as anxiety, depression, and intense craving during smoking abstinence as compared to men (Al’Absi, 2006; Leventhal et al., 2007; Panagiotakopoulos & Neigh, 2014; Pang et al., 2018; Perkins et al., 2012; Schnoll et al., 2007; Xu et al., 2008). The stress biomarker, cortisol is also higher in women than men during abstinence from smoking (Hogle & Curtin, 2006). In general, women exhibit lower smoking cessation rates, and are less likely to benefit from nicotine replacement therapy than men (Cepeda-Benito et al., 2004; Perkins 2001; Perkins & Scott, 2008; Piper et al., 2010). Although it is clear that women experience greater stress during smoking abstinence, the biological factors that promote nicotine withdrawal in females remain unclear.

A growing body of literature suggests that the ovarian hormones, estradiol (E2) and progesterone can influence the expression of negative affective states elicited during nicotine withdrawal. Specifically, women who quit in the luteal phase of the menstrual cycle experience greater negative affective states (distress scores and mood states) during withdrawal as compared to women who quit in the follicular phase (O’Hara et al., 1989; Perkins et al., 2000). A meta-analytic review also revealed that women report greater withdrawal during the luteal versus follicular phase (Weinberger et al., 2015). However, there are reports that do not find differences in withdrawal severity across women who quit during the follicular versus luteal phase of the menstrual cycle (Allen et al., 2000 & 2010). Although these studies suggest that ovarian hormones modulate nicotine withdrawal in women, the relationship between withdrawal severity and E2 and progesterone levels remains unclear.

Pre-clinical work has studied nicotine withdrawal in rodents following chronic exposure to nicotine. Following at least 7-14 days of nicotine exposure, withdrawal can be studied following the removal of the nicotine pump or following administration of a nicotinic receptor antagonist, such as mecamylamine (Kenny & Markou, 2001; Malin, 2001). The negative affective states produced by withdrawal elicit physical signs and anxiety-like behavior that can be assessed on the elevated plus maze (EPM) or light-dark-transfer (LDT) tests (Bruijnzeel, 2019; Jackson et al., 2015; Malin & Goyarzu, 2009; O’Dell & Khroyan, 2009). Previous rodent studies have assessed sex differences in the underlying biological factors that modulate nicotine withdrawal (Gentile et al., 2011; Hamilton et al., 2010; Kota et al., 2007 & 2008; Skwara et al., 2012; Torres et al., 2013 & 2015). Although these reports find sex differences during nicotine withdrawal, there are remaining questions with regard to the contribution of ovarian hormones, such as E2 and progesterone in the expression of anxiety-like behavior during withdrawal. Thus, the present study examined sex differences and the role of E2 and progesterone in the behavioral and biological indices of stress produced by nicotine withdrawal. *Study 1* compared nicotine withdrawal in male, intact female, and OVX rats. The magnitude of withdrawal was also compared in intact females that were tested in proestrus, estrus, metestrus, and diestrus. Lastly, the relationship between withdrawal severity and the levels of E2 and progesterone were assessed in intact females. *Study 2* examined whether ovarian hormone replacement influenced the expression of nicotine withdrawal in OVX rats that received vehicle, E2, or E2+progesterone supplementation.

## 2. Methods

### 2.1. Subjects

Male and female Wistar rats were bred in house from an out-bred stock of animals (Envigo, Inc.). On post-natal day (PND) 21, the rat pups were weaned and housed with a same-sex littermate for the remainder of the study. The rats were housed in a humidity- and temperature-controlled (22°C) vivarium on a reverse 12-hr light/dark cycle (lights off at 8:00 am and on at 8:00 pm) with *ad libitum* access to food and water. Prior to beginning the experiment, the rats were handled for 5 days. All the experimental procedures were approved by the UTEP Institutional Animal Care and Use Committee. The studies were conducted in compliance with the National Research Council, Guide for the Care and Use of Animals (8^th^ edition, 2010).

### 2.2. Overall design

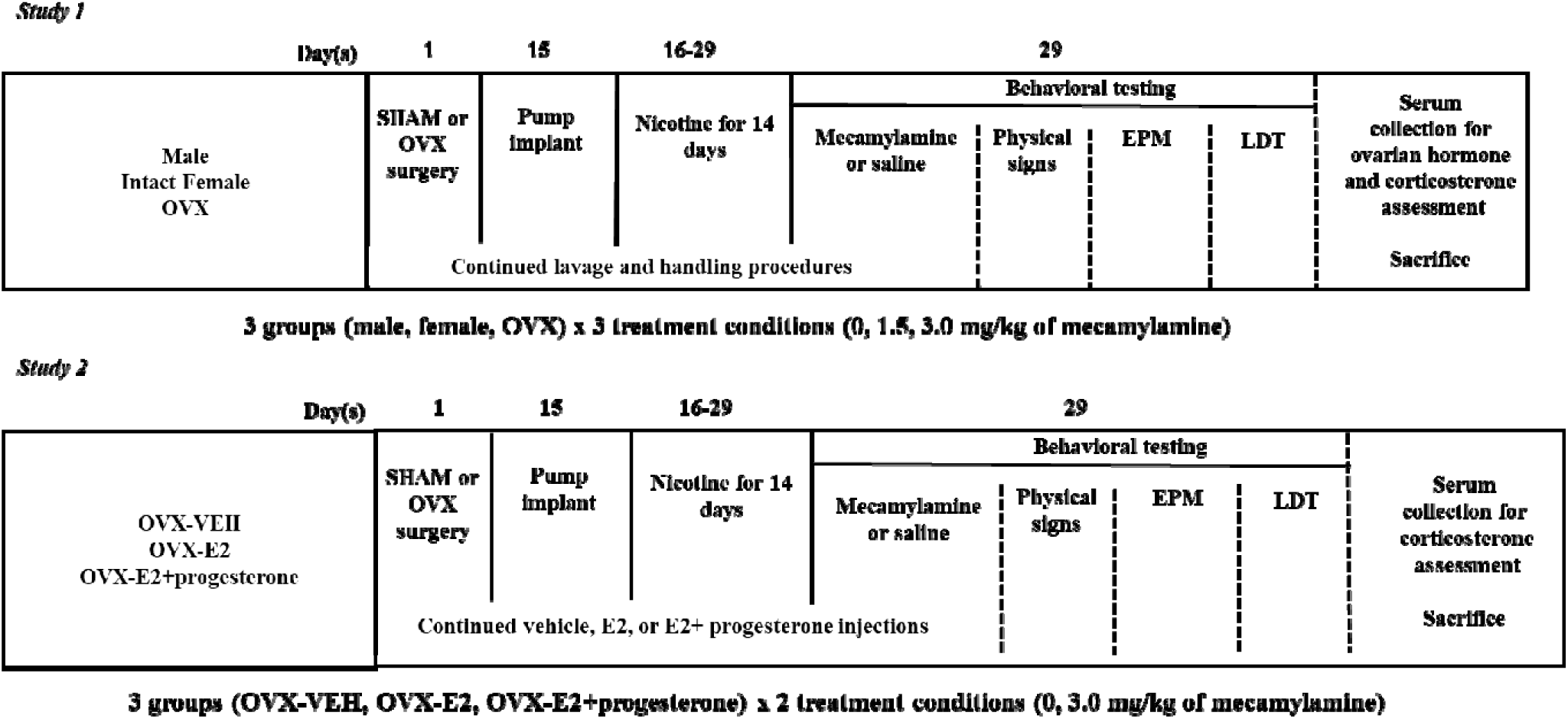

These diagrams depict our experimental groups and procedures. *Study 1* compared the magnitude of withdrawal in nicotine-dependent male, intact female, and OVX rats that received saline or mecamylamine (1.5 or 3.0 mg/kg) on the test day. *Study 2* also compared withdrawal in nicotine-dependent OVX rats that received vehicle, E2 or E2+progesterone immediately after the ovariectomy procedure. On the test day, the rats received saline or the highest dose of mecamylamine (3.0 mg/kg), since this dose elicited withdrawal in the OVX rats of Study 1. The rats were tested in a series of behavioral tests that included physical signs, EPM, and LDT tests. At the end of testing in Study 1, trunk blood was collected to assess serum E2 and progesterone levels. Also, the intact females received vaginal lavage procedures to assess the phase of the estrous cycle they were tested in. After behavioral testing in Study 1 and 2, trunk blood was collected to assess serum corticosterone levels.

### 2.3. Surgery

At PND 45-46, the rats were anesthetized using an isoflurane/oxygen mixture (1-3% isoflurane). Some of the females received surgical removal of ovarian tissue and the male and intact female rats received a sham procedure, as described in (Torres et al., 2009). The OVX was done at PND 45-46 because female rats that receive OVX procedures at PND 45 display a reduction in the rewarding effects of nicotine (Torres et al., 2009; Flores et al., 2016) and a suppression of anxiety-like behavior and stress-associated gene expression during nicotine withdrawal (Torres et al., 2013 & 2015). Once the rats reached PND 60, the rats were re-anesthetized and surgically prepared with an osmotic pump that delivered nicotine continuously for 14 days (3.2 mg/kg/day, expressed as base; model 2ML2; 5.0 uL/hour; Durect Corporation, Inc.). Previous studies have shown that this dose of nicotine produces similar levels of cotinine, the major nicotine metabolite in male and female Wistar adult rats (O’Dell et al., 2007).

### 2.4. Behavioral testing

The rats were placed into a clear Plexiglas^®^ cage in a test room that was dedicated to the assessments of physical signs of withdrawal under regular light conditions. Following a 10-min acclimation period, the rats received an injection of saline or mecamylamine (1.5 or 3.0 mg/kg). Ten min later, the physical signs of withdrawal were assessed for an additional 10 min, an observation period that has been used repeatedly in our laboratory (O’Dell et al., 2004; Tejeda et al., 2012). The observed signs included blinks, writhes, body shakes, teeth chatters, gasps, grooming bouts, and ptosis. Multiple successive counts of any sign required a distinct pause between episodes. Ptosis was only counted once per min. The total number of physical signs were defined as the sum of individual occurrences of the signs during the entire observation period. Following the physical signs assessment, the rats were transported to another dimly lit room for a 5 min acclimation period prior to the EPM test. The EPM apparatus consists of 4 arms (2 closed and 2 open) elevated to a height of 50 cm above the ground. The apparatus was illuminated by a red light suspended from the ceiling. At the beginning of the test, the rats were placed in the center of the EPM facing an open arm. Time spent in the center area, open, and closed arms was recorded for 5 min. Anxiety-like behavior was operationally defined as a decrease in time spent in the open arms relative to controls. Following EPM testing, the rats were transported to another room and acclimated for 5 min prior to the LDT test. The LDT apparatus consists of 2 enclosed chambers, one of which has clear Plexiglas^®^ walls and the other has solid black walls. The apparatus is separated by a wall with an opening that allowed the rats free access to both sides. The apparatus was positioned in the middle of the room under regular light conditions. At the start of the LDT test, the rats were placed in the middle of the dark chamber facing the back wall. Time spent in each side was recorded for 5 min. Anxiety-like behavior test was operationally defined as a decrease in time spent in the lit compartment relative to controls. The behavioral equipment was thoroughly cleaned and dried between each animal that was tested.

### 2.5. Hormone level assessments

After behavioral testing, the intact females in *Study 1* were sacrificed by rapid decapitation and trunk blood was collected. The blood was centrifuged for 15 min at 5000 x g at 4°C. Serum was extracted and stored in 100 µL aliquots at −80°C until analyzed via enzyme-linked immunosorbent assay (ELISA) procedures for progesterone (Enzo Life Sciences, Farmingdale, USA) and corticosterone (Assaypro, Winfield, MO), according to the manufacturer instructions. Standards with known concentrations were included in each assay, and they ranged from 0 to 100 ng/mL for corticosterone and 0 to 500 pg/mL for progesterone. Samples were placed in a 96 well-plate and read at 450 and 630 nm wavelength using a Spectra Maxplus spectrophotometer (Molecular Devices, Inc.). E2 levels were assayed at the University of Pittsburgh Small Molecule Biomarker Core, using liquid chromatography-tandem mass spectrometry (LC-MS/MS). LC-MS/MS is the preferred method to estimate E2 levels given that the serum concentration of this hormone is below a detectable range of sensitivity for standard immunoassays, such as ELISA (Field, 2013).

### 2.6. Estrous cycle determination

In *Study 1*, vaginal lavage and cytology procedures were conducted in intact females in order to determine proestrus, estrus, metestrus, or diestrus, as described previously (see Goldman et al., 2007; Torres et al., 2009). The lavage procedures began 8 days prior to the pump surgery and continued until the end of the test day. In *Study 2*, the OVX rats received lavage procedures only on the test day. The OVX rats that received vehicle displayed a metestrus cytology, E2 supplementation a proestrus cytology, and E2+progesterone an estrus cytology (data not shown).

### 2.7. Hormone supplementation procedures

*In Study 2*, OVX rats received a 4-day hormone supplementation procedure that began the day after the ovariectomy surgery. OVX control rats received 3 repeated vehicle injections (peanut oil, sc) and no injection on the 4^th^ day. To examine the effects of E2 alone, a group of OVX rats received a 0.1 mL bolus injection of E2 (5 µg, sc) for the first 2 days, a vehicle injection on the 3^rd^ day, and no injection on the 4th day. This supplementation procedure mimics normal E2 cycling patterns in intact female rats (see Asarian et al., 2002). On the test day, rats received their E2 injection 30 min before the behavioral battery. To examine the effects of E2 and progesterone, a group of OVX rats received a 0.1 mL bolus injection of E2 (5 µg, sc) for the first 2 days, a 0.1 mL bolus injection of progesterone (250 µg, sc) on the 3^rd^ day, and no injection on the 4^th^ day. On the test day, rats received their progesterone injection 4.5 hours before the behavioral battery. The 4-day supplementation procedure was repeated 5 times prior to testing. A group that received progesterone alone was not included because progesterone only induces sexual receptivity in OVX rats that are supplemented with E2+progesterone (Becker et al., 2005).

### 2.8. Statistics

In *Study 1*, the data were analyzed separately for physical signs, EPM, LDT, and corticosterone levels using 2-way analysis of variance (ANOVA) with group (male, intact female, and OVX) and treatment condition (0, 1.5, 3.0 mg/kg of mecamylamine) as between subjects factors. Where appropriate, significant interaction effects were further analyzed using post-hoc comparisons (Fisher’s LSD test, *p*≤0.05). To examine estrous cycle effects, the intact females that received the 3.0 mg/kg dose of mecamylamine were grouped according to the estrous phase they were tested in. The data were then analyzed separately for physical signs, EPM, LDT, and corticosterone levels using a 1-way ANOVA with stage of estrous (proestrus, estrus, metestrus, diestrus) as a between subjects factor. The relationship between hormones and behavior was then examined via simple linear regression. Each hormone was correlated separately with each measure of withdrawal. A Pearson correlation co-efficient was computed to assess the strength and direction of each relationship. A 1-sample t-test was used to examine whether the correlations were statistically significant. In intact female rats, the relative concentration of E2 and progesterone varies across the estrous cycle. Therefore, a statistical approach was employed to assess whether female rats could be categorized into groups that displayed similar concentrations of these hormones. A k-means clustering algorithm was used to classify rats according to their levels of E2 and progesterone. This analysis resulted in 2 groups that had significantly different hormone levels (high E2/low progesterone and low E2/high progesterone). Individual rats were grouped and differences in physical signs, EPM, LDT, and corticosterone levels were compared using a between subjects t-test. This k-means categorization process has been used to classify older and younger animals according to their performance in working memory and behavioral flexibility tasks (Mota et al., 2019).

In *Study 2*, the data were analyzed separately for physical signs, EPM, LDT, and corticosterone levels using 2-way ANOVA with group (OVX-veh, OVX-E2, OVX-E2+progesterone) and treatment (0, 3.0 mg/kg of mecamylamine) as between subjects factors. Where appropriate, significant interaction effects were further analyzed using post-hoc comparisons (Fisher’s LSD test, *p*≤0.05).

## 3. Results

### 3.1. Study 1

*Figure 1* displays physical signs (A), EPM (B), LDT (C), and corticosterone levels (D) in nicotine-treated male, intact female, and OVX rats that received mecamylamine (0, 1.5, or 3.0 mg/kg) on the test day. With regard to physical signs, there was no interaction between group and treatment conditions [F(4,154)=0.57; *p*=0.58]. However, there was a main effect of treatment condition [F(2,154)=98.33; *p*≤0.05], with all rats that received mecamylamine displaying more physical signs as compared to rats that received saline (\**p*≤0.05).

**Figure 1.**
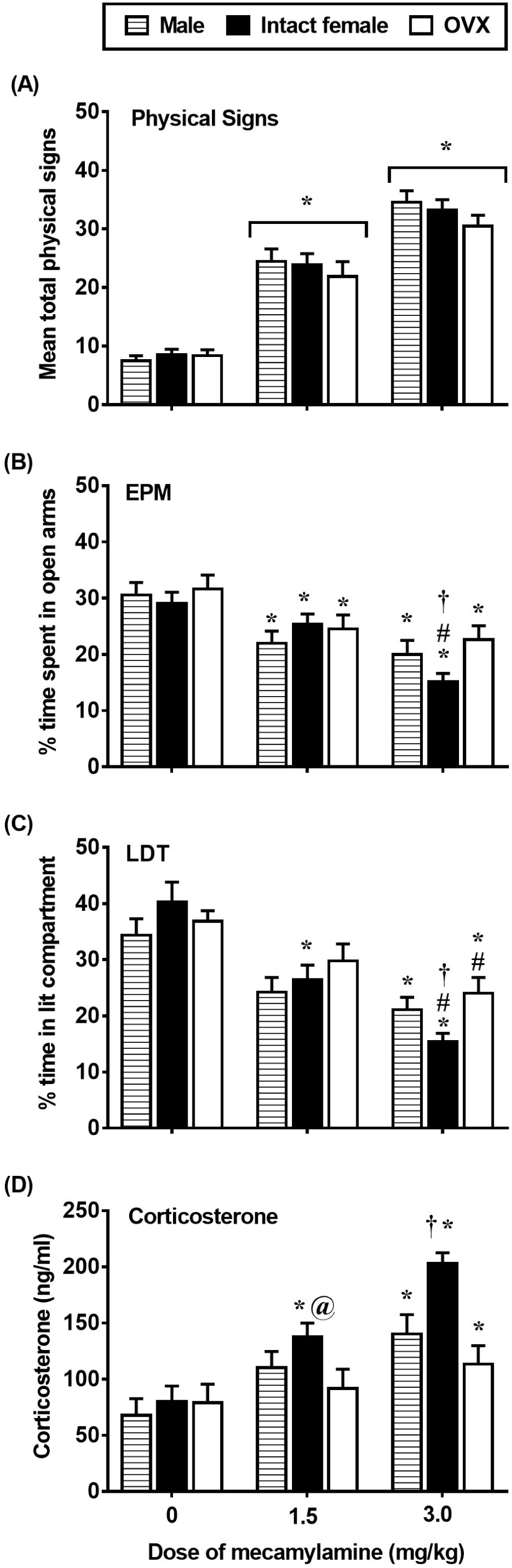
Mean (±SEM) physical signs (A), % time in the open arms (B), % time in the lit compartment (C), and corticosterone levels (D) in male (0 n=16; 1.5 n=17; 3.0 n=12), intact female (0 n=18; 1.5 n=24; 3.0 n=38), and OVX (0 n=13; 1.5 n=12; 3.0 n=13) rats. Asterisks (*) denote a difference from respective saline controls, number signs (#) denote a difference from their respective mecamylamine-treated group, at signs (@) denote a difference from OVX rats in their respective treatment condition, and the dagger (†) denotes a difference from males and OVX rats in their respective treatment condition (*p*≤0.05).

With regard to the EPM data, a 2-way ANOVA of % time in open arms revealed a significant interaction between group and treatment condition [F(4,154)= 2.87; *p*≤0.05]. In males, post-hoc analyses revealed a significant decrease in % time in open arms in rats that received the 1.5 or 3.0 mg/kg dose of mecamylamine as compared to saline controls (\**p*≤0.05). There was no difference between males that received the 1.5 or 3.0 mg/kg dose of mecamylamine (*p*=0.47). In intact females, post-hoc analyses revealed a significant decrease in % time in open arms in rats that received the 1.5 or 3.0 mg/kg dose of mecamylamine as compared to saline controls (\**p*≤0.05). The % time in open arms was lower in intact females that received the 3.0 versus 1.5 mg/kg dose of mecamylamine (#*p*≤0.05). In OVX rats, post-hoc analyses revealed a significant decrease in % time in open arms in rats that received the 1.5 or 3.0 mg/kg dose of mecamylamine as compared to their respective saline controls (\**p*≤0.05). There was no difference between OVX rats that received the 1.5 versus 3.0 mg/kg dose of mecamylamine (\**p*=0.59). Group differences were detected in rats that were treated with the 3.0 mg/kg dose of mecamylamine. Post-hoc analyses revealed that % time in open arms was lower in intact females as compared to male and OVX rats (†*p*≤0.05). An analysis of closed arm entries was conducted to examine whether our assessment of anxiety-like behavior was influenced by group differences in locomotor activity (see *Table 2*). The analysis revealed that there were no group or treatment effects in closed arm entries [F (4, 154)= 0.09, *p*=0.98].

**Table 1.**
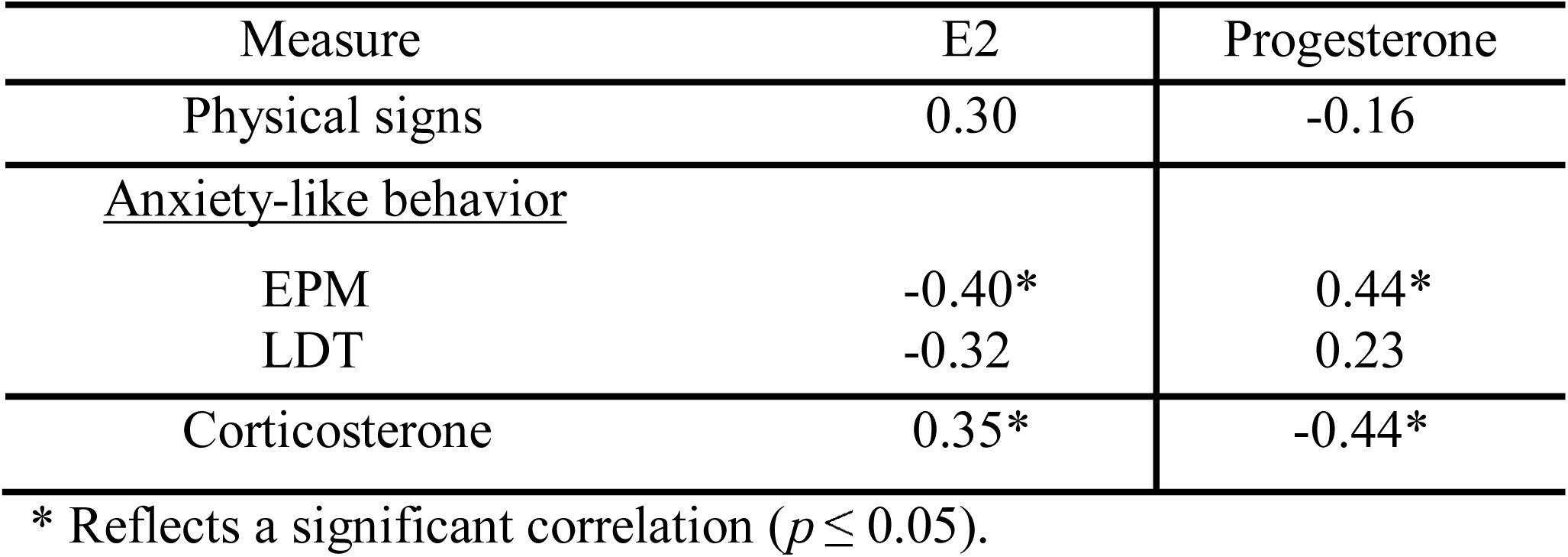
Correlation values (Pearson’s r) between behavioral measures of nicotine withdrawal or corticosterone levels with ovarian hormone levels

**Table 2.** Closed arm entries in the EPM (mean ± SEM)

| Study 1 | Group: | Male | Female | OVX |
| --- | --- | --- | --- | --- |
| | Saline | 10.37 ± 0.75 | 10.94 ± 0.71 | 11.31 ± 0.84 |
| | Mecamylamine 3.0 mg/kg | 10.23 ± 0.73 | 10.54 ± 0.61 | 11.09 ± 0.87 |
| | Mecamylamine 3.0 mg/kg | 10.33 ± 0.87 | 10.34 ± 0.49 | 10.39 ± 0.83 |
| Study 2 | Group: | OVX-veh | OVX-E2 | OVX-E2+progesterone |
| | Saline | 11.50 ± 0.63 | 10.23 ± 0.55 | 10.53 ± 0.55 |
| | Mecamylamine 3.0 mg/kg | 10.50 ± 0.71 | 10.90 ± 0.63 | 10.23 ± 0.55 |

With regard to the LDT data, a 2-way ANOVA of % time the lit compartment revealed a significant interaction between group and treatment condition [F(4,154)=3.02; *p*≤0.05]. In males, post-hoc analyses revealed a significant decrease in % time in the lit compartment in rats that received the 3.0 (\**p*≤0.05), but not 1.5 (*p*=0.06) mg/kg dose of mecamylamine as compared to saline controls. There was no difference between males that received the 1.5 or 3.0 mg/kg dose of mecamylamine (*p*=0.25). In intact females, post-hoc analyses revealed a significant decrease in % time in the lit compartment in rats that received the 1.5 or 3.0 mg/kg dose of mecamylamine as compared to their respective saline controls (\**p*≤0.05). The % time in the lit compartment was lower in intact females that received the 3.0 versus 1.5 dose mg/kg of mecamylamine (#*p*≤0.05). In OVX rats, post-hoc analyses revealed a significant decrease in % time in the lit compartment in rats that received the 3.0 (\**p*≤0.05), but not 1.5 (*p*=0.28) mg/kg dose of mecamylamine as compared to their respective saline controls. The % time in the lit compartment was lower in OVX rats that received the 3.0 versus 1.5 mg/kg dose of mecamylamine (#*p*≤0.05). Group differences were detected in rats that were treated with the 1.5 mg/kg dose of mecamylamine. Post-hoc analyses revealed that % time in the lit compartment was lower in intact females as compared to OVX rats (@*p*≤0.05). Group differences were also detected in rats that were treated with the 3.0 mg/kg dose of mecamylamine. Post-hoc analyses revealed that % time in the lit compartment was lower in intact females as compared to male and OVX rats (†*p*≤0.05).

With regard to corticosterone levels, a 2-way ANOVA revealed a significant interaction between group and treatment condition [F(4,154)=2.60; *p*≤0.05]. In males, post-hoc analyses revealed a significant decrease in % time in the lit compartment in rats that received the 1.5 or 3.0 mg/kg dose of mecamylamine as compared to their respective saline controls (\**p*≤0.05). There was no difference between males that received the 1.5 or 3.0 mg/kg dose of mecamylamine (*p*=0.18). In intact females, post-hoc analyses revealed a significant increase in corticosterone levels in rats that were treated with the 1.5 or 3.0 mg/kg dose of mecamylamine as compared to their respective saline controls (\**p*≤0.05). Corticosterone levels were also higher in intact females that received the 3.0 versus 1.5 mg/kg dose of mecamylamine (*p*≤0.05). In OVX rats, post-hoc analyses revealed a significant increase in corticosterone levels in rats that were treated with the 3.0 (\**p*≤0.05), but not 1.5 (*p*=0.59) mg/kg dose of mecamylamine as compared to saline controls. Corticosterone levels did not differ in OVX rats that received the 1.5 versus the 3.0 mg/kg dose of mecamylamine (*p*=0.34). Group differences were detected across rats that were treated with the 1.5 mg/kg dose of mecamylamine. Specifically, post-hoc analyses revealed that corticosterone levels were higher in intact females as compared to OVX rats (@*p*≤0.05); however, there was no difference between intact female and male rats (*p*=0.41). Group differences were detected across rats that were treated with the 3.0 mg/kg dose of mecamylamine. Post-hoc analyses revealed that corticosterone levels were higher in intact females as compared to both male and OVX rats (†*p*≤0.05).

*Figure 2* displays E2 (A) and progesterone (B) levels in intact females that received mecamylamine (3.0 mg/kg) and were tested during various phases of the estrous cycle. With regard to E2, a 1-way ANOVA revealed a significant interaction between group and treatment condition [F(3,34)=3.63; *p*≤0.05]. Post-hoc analyses revealed that females that were tested in estrus displayed lower E2 as compared to females in all other phases combined (†*p*≤0.05). With regard to progesterone, a 1-way ANOVA revealed a significant interaction between group and treatment condition [F(3,34)=12.20; *p*≤0.05]. Post-hoc analyses revealed that females that were tested in estrus displayed higher progesterone levels as compared to rats that were tested in all other phases combined (†*p*≤0.05).

**Figure 2.**
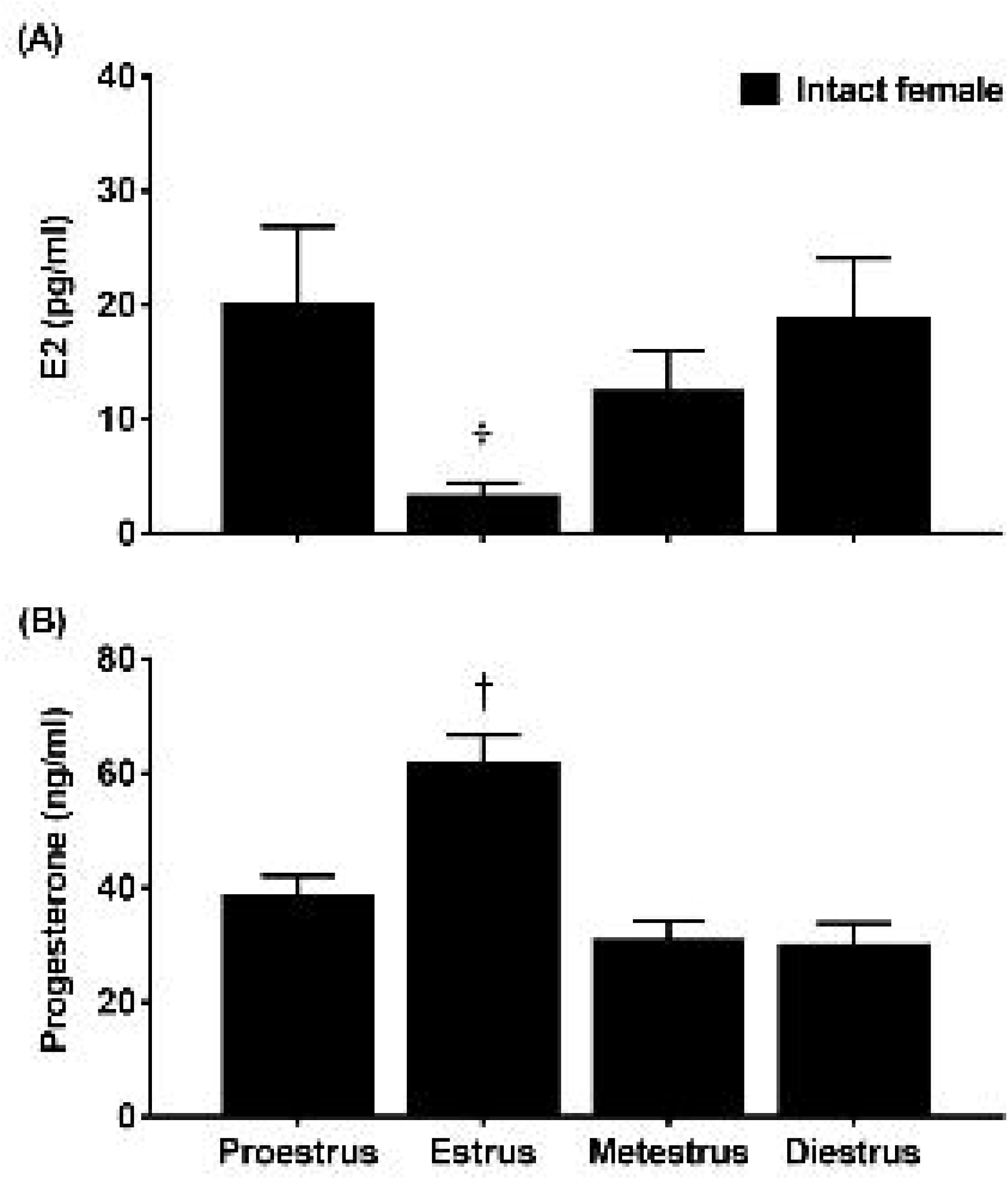
Mean (±SEM) E2 (A) and progesterone (B) levels following administration of mecamylamine (3.0 mg/kg) in nicotine-treated intact females that were tested in proestrus (n=12), estrus (n=12), metestrus (n=7), or diestrus (n=7). Daggers (†) denote a significant difference from all other groups (*p*≤0.05).

*Figure 3* displays physical signs (A), EPM (B), LDT (C), and corticosterone levels (D) in intact females that received mecamylamine (3.0 mg/kg) and were tested during various phases of the estrous cycle. With regard to physical signs, a 1-way ANOVA revealed that there was no main effect of estrous cycle [F(3, 34) = 0.42; *p*=0.74].With regard to the EPM data, a 1-way ANOVA of % time in open arms revealed a main effect of estrous cycle [F(3, 34)=4.96; *p*≤0.05]. Post-hoc analyses revealed that females that were tested in estrus displayed greater % time in the open arms as compared to females that were tested in proestrus, metestrus, or diestrus (†*p*≤0.05). With regard to the LDT data, a 1-way ANOVA of % time in the lit compartment revealed a main effect of estrous cycle [F(3, 34)=3.14; *p*≤0.05]. Post-hoc analyses revealed that females that were tested in estrus displayed greater % time in the lit compartment as compared to females in proestrus, metestrus, or diestrus (†*p*≤0.05). With regard to corticosterone, a 1-way ANOVA revealed a main effect of estrous cycle [F(3, 34)=3.11; *p*≤0.05]. Post-hoc analyses revealed that females that were tested in estrus displayed lower corticosterone levels as compared to all other phases combined (†*p*≤0.05).

**Figure 3.**
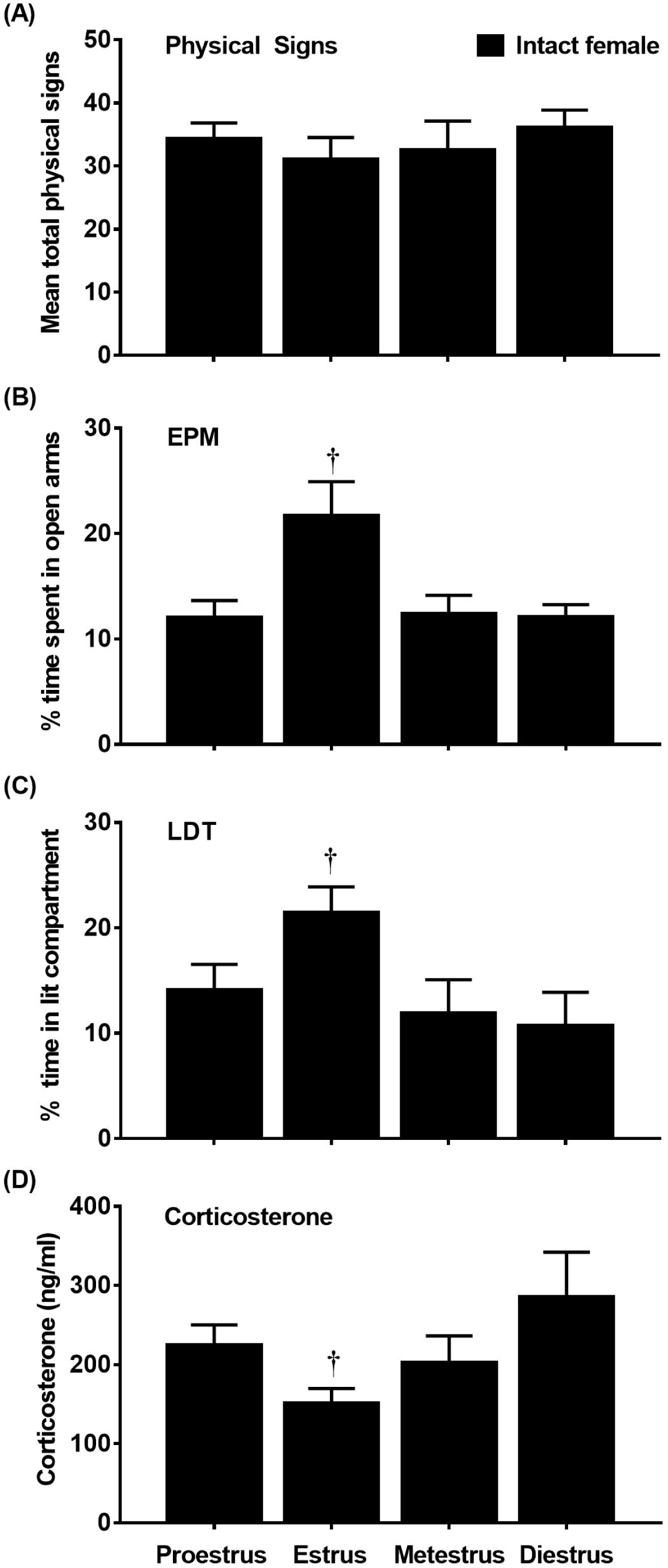
Mean (±SEM) physical signs (A), % time in the open arms (B), % time in the lit compartment (C) and corticosterone (D) levels following administration of mecamylamine (3.0 mg/kg) in nicotine-treated intact females that were tested in proestrus (n=12), estrus (n=12), metestrus (n=7), or diestrus (n=7). Daggers (†) denotes a difference from all other groups (*p*≤0.05).

*Table 1* displays correlation values (Pearson’s r) between behavioral measures of withdrawal or corticosterone levels with E2 or progesterone levels. With regard E2, there was no correlation between E2 and physical signs of withdrawal (r=0.29, *p*=0.07). However, there was a negative correlation between E2 and % time in open arms of the EPM (r=-0.38; *p*≤0.05). There was a negative correlation between E2 and % time in the lit compartment of the LDT (r=-0.32; *p*≤0.05). There was a positive correlation between E2 and corticosterone levels (r=0.35; *p*≤0.05). With regard to progesterone, there was no correlation between progesterone and physical signs of withdrawal (r=-0.16; *p=*0.35). There was a positive correlation between progesterone and % time in open arms of the EPM (r=0.44; *p*≤0.05); however, there was no correlation between progesterone and % time in the lit compartment of the LDT (r=0.22; *p*=0.26). There was a negative correlation between progesterone and corticosterone levels (r=-0.43; *p*≤0.05).

*Figure 4* displays data from the 2 groups of intact females that displayed either high E2/low progesterone or low E2/high progesterone levels. The graph depicts E2 and progesterone levels (A), physical signs (B), EPM (C), LDT (D), and corticosterone levels (E). The cluster analysis revealed 2 groups of intact females that displayed high E2/low progesterone or low E2/high progesterone levels. With regard to E2, there was a significant difference between the high E2/low progesterone and the low E2/high progesterone group (t(36)=3.17, *p*≤0.05). With regard to progesterone, there was a significant difference between the low E2/high progesterone and the high E2/low progesterone group (t(36)=16.2, *p*≤0.05). With regard to physical signs, there were no differences between the high E2/low progesterone versus low E2/high progesterone groups (t(36)=1.14, *p*=0.26). With regard to the EPM data, the group that displayed high E2/low progesterone displayed less % time in the open arms of the EPM as compared to females that displayed low E2/high progesterone (t(36)=3.50; *p*≤0.05). With regard to the LDT data, the group that displayed high E2/low progesterone spent less time in the lit compartment as compared to the group that displayed low E2/high progesterone (t(36)=3.24, *p*≤0.05). With regard to corticosterone, the group that displayed high E2/low progesterone displayed higher corticosterone levels as compared to the group that displayed low E2/high progesterone (t(36)=2.12, *p*≤0.05).

**Figure 4.**
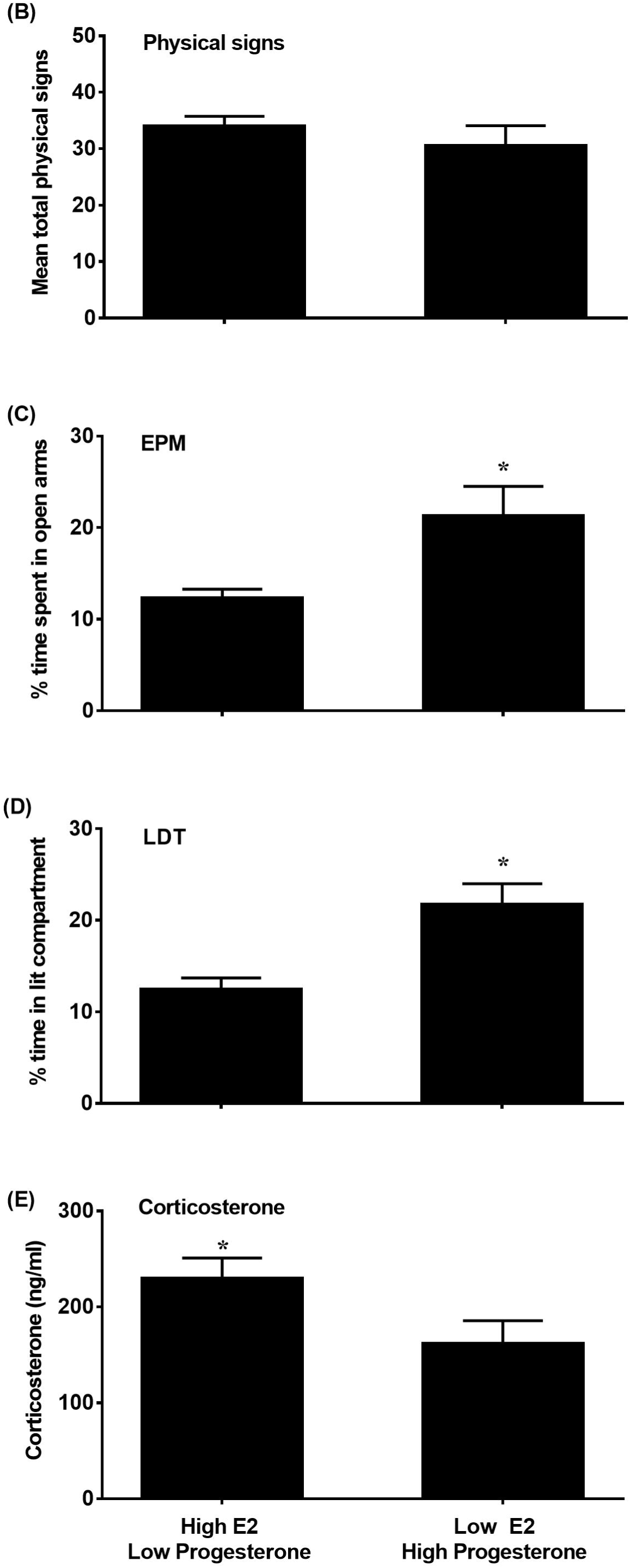
Mean (±SEM) E2 and progesterone levels (A), physical signs (B), % time in the open arms (C), % time in the lit compartment (D) and corticosterone levels (E) following administration of mecamylamine (3.0 mg/kg) in nicotine-treated intact females that displayed high E2/low progesterone (n=26) or low E2/high progesterone (n=12) levels.

### 3.2. Study 2

*Figure 5* displays physical signs (A), EPM (B), LDT (C), and corticosterone levels (D) in OVX rats that received vehicle (OVX-vehicle), E2 (OVX-E2), or both E2 and progesterone (OVX-E2+progesterone). With regard to physical signs, there was no interaction between group and treatment condition [F(2,61)=1.96; *p*=0.15]. However, there was a main effect of treatment condition [F(1,61)=441.95; *p*≤0.05], with rats that received the 3.0 mg/kg dose of mecamylamine displaying more physical signs as compared to saline controls (\**p*≤0.05). There were no differences in withdrawal signs across groups.

**Figure 5.**
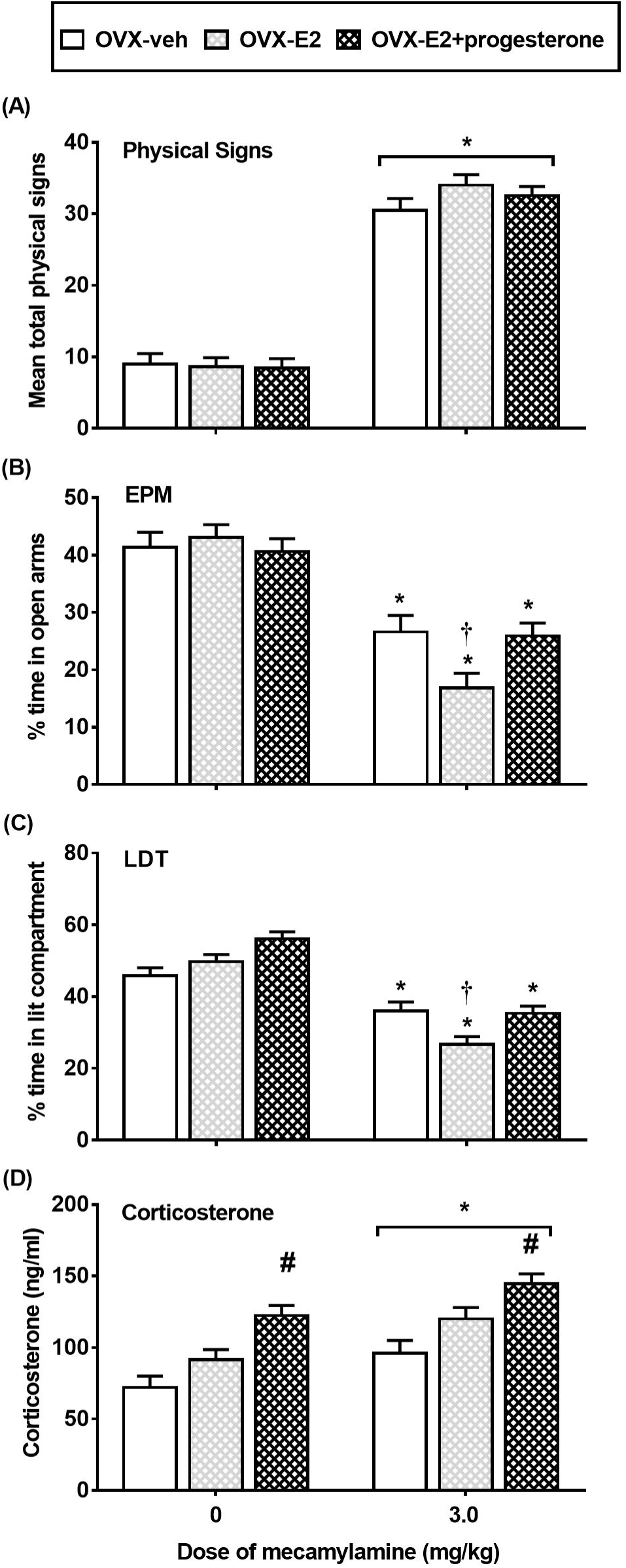
Mean (±SEM) physical signs (A), % time in the open arms of the EPM (B), % time in the lit compartment (C) and corticosterone levels (D) in nicotine-treated OVX rats that received vehicle (OVX-veh 0 n=10; 3.0 n=8), E2 (OVX-E2; 0 n=13; 3.0 n=10) or E2+progesterone (OVX-E2+progesterone; 0 n=13; 3.0 n=13). Asterisks (*) denote a difference from saline controls, number signs (#) denote a difference from OVX-veh and OVX-E2 regardless of mecamylamine treatment, and the daggers (†) denote a difference from OVX-veh and OVX-E+progesterone rats in their respective treatment condition (*p*≤0.05).

With regard to the EPM data, a 2-way ANOVA revealed a significant interaction between group and treatment condition [F(2,61)=3.70; *p*≤0.05]. Post-hoc analyses revealed that % time in open arms was lower in rats that received the 3.0 mg/kg dose of mecamylamine as compared to saline controls, an effect that was significant in OVX-vehicle, OVX-E2, and OVX-E2+progesterone rats (\**p*≤0.05). Group differences were detected in OVX rats that received the 3.0 mg/kg dose of mecamylamine. Post-hoc analyses revealed that % time in open arms was lower in OVX-E2 rats as compared to OVX-vehicle and OVX-E2+progesterone (†*p*≤0.05) rats. *Table 2* displays our analysis of closed arm entries. The analysis revealed that there were no group or treatment effects in closed arm entries [F (2, 61)= 0.90, *p* = 0.41].

With regard to the LDT data, a 2-way ANOVA revealed a significant interaction between group and treatment condition [F(2,61)=4.82; *p*≤0.05]. Post-hoc analyses revealed a significant decrease in % time in the lit compartment in rats that received the 3.0 mg/kg dose of mecamylamine as compared to saline controls, an effect that was significant in OVX-vehicle, OVX-E2, and OVX-E2+progesterone rats (\**p*≤0.05). Group differences were detected in rats that received the 3.0 mg/kg dose of mecamylamine. Post-hoc analyses revealed that % time in the lit compartment was lower in OVX-E2 rats as compared to OVX-vehicle and OVX-E2+progesterone rats (†*p*≤0.05).

With regard to corticosterone, a 2-way ANOVA revealed that there was no interaction between group and treatment condition [F(2, 61)=0.91; *p*=0.12]. However, there was a main effect of treatment condition [F(1,61)=15.56; *p*≤0.05], with rats that received the 3.0 mg/kg dose of mecamylamine displaying higher corticosterone levels as compared to saline controls (\**p*≤0.05). Also, there was a main effect of group [F(1,61)=20.80; *p*≤0.05]. Regardless of treatment condition, OVX-E2+progesterone rats displayed higher corticosterone levels as compared to OVX-vehicle and OVX-E2 rats (#*p*≤0.05).

## 4. Discussion

The major finding of this report is that intact female rats displayed greater indices of stress produced by nicotine withdrawal as compared to males, and this effect was ovarian hormone-dependent. In *Study 1*, intact females displayed higher anxiety-like behavior and corticosterone levels during withdrawal as compared to male and OVX rats. With regard to the estrous cycle, intact females that were tested in estrus (when E2 levels are relatively low) displayed less anxiety-like behavior and corticosterone levels as compared to all other phases. In intact females, the magnitude of anxiety-like behavior and corticosterone levels were positively correlated with E2 and negatively correlated with progesterone levels. In intact female rats, 2 groups emerged from a cluster analysis that displayed either high E2/low progesterone or low E2/high progesterone levels. Females displaying high E2/low progesterone displayed greater anxiety-like behavior and corticosterone levels during withdrawal as compared to the other group. In *Study 2*, OVX rats that received E2 supplementation displayed greater anxiety-like behavior as compared to OVX rats that received vehicle or E2+progesterone. All groups of OVX rats displayed an increase in corticosterone levels during withdrawal regardless of the supplementation condition.

The present study found that there were no sex differences in the physical signs of withdrawal, consistent with previous studies from our laboratory (Torres et al., 2013; Correa et al., 2019) and others using similar lighting conditions (Hamilton et al., 2010). Also, the physical signs of withdrawal do not appear to be influenced by ovarian hormones, as OVX rats displayed similar physical signs as compared to male and intact female rats. Also, the physical signs of withdrawal were similar across female rats that were tested in different phases of the estrous cycle. This finding is not consistent with a recent report showing that physical signs produced by nicotine withdrawal were higher in female rats that were tested in metestrus versus proestrus (Henceroth et al., 2018). The physical signs of withdrawal were also similar in intact females that displayed either high E2/low progesterone or low E2/high progesterone levels. Lastly, the physical signs of withdrawal were not correlated with E2 and progesterone levels and they were similar across groups of OVX rats that received vehicle, E2, and E2+progesterone supplementation. Together, these data suggest that ovarian hormones *do not* play a role in modulating physical signs produced by nicotine withdrawal.

With regard to sex differences, a major finding of this report is that intact females displayed greater anxiety-like behavior and corticosterone levels during nicotine withdrawal than males. These findings are consistent with previous work showing that during nicotine withdrawal, female rats display higher levels of anxiety-like behavior than males (Torres et al., 2013). Reports from other laboratories have also revealed that during nicotine withdrawal, female rats display higher serum levels of corticosterone and adrenocorticotropic hormone than males (Gentile et al., 2011; Skwara et al., 2012). The notion that female rodents experience greater negative affective states during withdrawal is consistent with the finding that both female rats and mice display greater place aversion to an environment paired previously with nicotine withdrawal (Kota et al., 2007 & 2008; O’Dell & Torres, 2014).

Through a series of different approaches, *Study 1* revealed that ovarian hormones play a major role in stress responses produced by nicotine withdrawal. <u>First</u>, OVX females displayed less anxiety-like behavior and corticosterone levels as compared to intact females, consistent with previous work (Torres et al., 2015). These studies suggest that the presence of ovarian hormones is important for the expression of stress responses elicited by nicotine withdrawal. <u>Second</u>, females that were tested in estrus (when E2 levels are relatively low) displayed less anxiety-like behavior and corticosterone levels as compared to all other phases. A previous report revealed that estrus females display less anxiety-like behavior in the EPM following presentation of a social stressor as compared to diestrus females (McCormick et al., 2008). Shansky et al. (2004) also found that estrus females displayed less impairments in a learning task following administration of a pharmacological stressor as compared to proestrus females. Other studies have also reported that the increase in corticosterone levels produced by presentation of an acute stressor was lower in estrus versus proestrus female rats (Viau & Meaney 1991; Conrad et al., 2004). Baseline measurements of anxiety-like behavior have also been shown to be lower in estrus versus diestrus females (Frye & Rhodes et al., 2006; Gouveia et al., 2004; Marcondes et al., 2001; Mora et al., 1996). Also, corticosterone levels are lower in estrus versus proestrus females following acute restraint stress (Figueiredo et al., 2002). These studies suggest that stress responses are lower during estrus, when E2 is lower and progesterone is higher relative to all other phases of the estrous cycle (Bohler., et al., 1990; Nappi et al., 1997). <u>Third</u>, anxiety-like behavior on the EPM and corticosterone levels were positivity correlated with E2 and negatively correlated with progesterone levels. Consistent with this finding, the cluster analysis revealed that rats displaying high E2/low progesterone displayed greater anxiety-like behavior in the EPM and LDT tests. Together, these findings suggest that E2 promotes and progesterone reduces the magnitude of anxiety-like behavior and corticosterone levels during nicotine withdrawal.

As another approach to study the role of ovarian hormones, *Study 2* assessed nicotine withdrawal in OVX rats that received E2 or E2+progesterone supplementation. The results revealed that OVX rats that received E2 displayed higher levels of anxiety-like behavior as compared to OVX rats that received vehicle. This finding is consistent with our assertion that E2 promotes anxiety-like behavior produced by nicotine withdrawal. The ability of E2 to increase sensitivity to a stressful stimulus in females may be due to the direct effects of E2 on the hypothalamus pituitary adrenal axis. This is based on our finding that OVX rats displayed reduced basal corticosterone levels, an effect that was reversed by E2 supplementation. Also, previous work revealed that the increase in corticosterone levels produced by acute stress are greater in OVX rats that received E2 supplementation (Handa & Weiser, 2014; Green et al., 2018). Also, the highest levels of the stress hormone, corticotropin releasing factor (CRF) have been observed during proestrus, when E2 levels are highest (Bohler et al., 1990; Nappi et al., 1997). The present study also revealed that the OVX-E2+progesterone rats displayed less anxiety-like behavior as compared to the OVX-E2 rats. This might have been related to the direct anxiolytic effects of progesterone that was administered prior to the test. Indeed, previous work has shown that progesterone administration decreases anxiety-like behavior in intact female and OVX mice (Mora et al., 1996; Reddy et al., 2005). These data suggest that progesterone may decrease anxiety-like behavior produced by nicotine withdrawal in females.

The present work is significant because it reflects an important first step towards understanding the role of ovarian hormones in modulating tobacco use in females. This report revealed that E2 promotes whereas progesterone reduces anxiety-like behavior produced by nicotine withdrawal. These results may inform the development of more effective tobacco cessation strategies in females. For example, a medical professional might assess the hormone status of a woman who is contemplating quitting smoking. Our data suggest that the best time to quit smoking may be in phases of the cycle when E2 levels are low and progesterone levels are relatively high in order to minimize the extent to which E2 may intensify the nicotine withdrawal syndrome. Indeed, a clinical report revealed that women have an easier time quitting smoking during the luteal phase, when E2 levels are decreasing and progesterone levels are relatively higher than E2 (Allen et al., 2008). During acute smoking abstinence, high levels of progesterone were associated with reduced negative affective states in nicotine-dependent women (Pang et al., 2018). Lastly, Nakajima et al. (2019) found reduced cortisol responses to stressful stimuli in the luteal versus the follicular phase in a sample of nicotine-dependent women. Future studies are needed to better understand the underlying biological factors that modulate the aversive effects nicotine withdrawal and promote tobacco use in females. This work will be an essential step towards developing more effective cessation strategies that will reduce health disparities produced by tobacco use in women.

## Acknowledgements

This research was supported by the National Institutes of Health (NIH; R01-DA021274 and R25-DA033613). Rodolfo J. Flores was a fellow in the Interdisciplinary Research Training Institute (R25-DA026401), Kevin P. Uribe was funded by a Ruth L. Kirschstein pre-doctoral fellowship (F31-DA046126), and Dr. Victor L. Correa was funded by a post-doctoral training contract from the National Institute on Drug Abuse (HHSN271201600057C). The authors thank Melissa Ibarra, Paola Correa, and Grace Hendricks for their technical assistance.

The authors declare no conflict of interest.

